# Inflammaging in human photoexposed skin: Early onset of senescence and imbalanced epidermal homeostasis across the decades

**DOI:** 10.1101/2022.03.28.486066

**Authors:** Bradley B. Jarrold, Christina Yan Ru Tan, Chin Yee Ho, Ai Ling Soon, TuKiet T. Lam, Xiaojing Yang, Calvin Nguyen, Wei Guo, Yap Ching Chew, Yvonne M. DeAngelis, Lydia Costello, Paola De Los Santos Gomez, Stefan Przyborski, Sophie Bellanger, Oliver Dreesen, Alexa B. Kimball, John E. Oblong

## Abstract

Inflammaging is a theory of aging which purports that low-level chronic inflammation leads to cellular dysfunction and premature aging of surrounding tissue. Skin is susceptible to inflammaging because it is the first line of defense from the environment, particularly solar radiation. To better understand the impact of aging and photoexposure on epidermal biology we performed a systems biology-based analysis of photoexposed face and arm and photoprotected buttock sites from women between the ages of 20’s to 70’s. Biopsies were analyzed by histology, transcriptomics, and proteomics and skin surface biomarkers collected from tape strips. We identified morphological changes with age of epidermal thinning, rete ridge pathlength loss, and stratum corneum thickening. The SASP biomarkers IL-8 and IL-1RA/IL1-α were consistently elevated in face across age and *cis/trans*-urocanic acid were elevated in arms and face with age. In older arms, the DNA damage response biomarker 53BP1 showed higher puncti numbers in basal layers and epigenetic aging was accelerated. Genes associated with differentiation and senescence show increasing expression in the 30’s whereas genes associated with hypoxia and glycolysis increase in the 50’s. Proteomics comparing 60’s vs 20’s confirmed elevated levels of differentiation and glycolytic related proteins. Representative immunostaining for proteins of differentiation, senescence, and oxygen sensing/hypoxia shows similar relationships. This systems biology-based analysis provides a body of evidence that young photoexposed skin is undergoing inflammaging. We propose the presence of chronic inflammation in young skin contributes to an imbalance of epidermal homeostasis that leads to a prematurely aged appearance during later life.

## 1 INTRODUCTION

The skin is the largest organ of the human body, providing protection from external insults such as solar radiation, pollution, chemicals, and particulate matter. Like all organs of the body the skin is susceptible to aging, resulting in structural and functional changes which may be accelerated further by environmental insults.^1^ This premature aging of skin leads to cellular and morphological changes that accumulate over time and ultimately affect the skin’s appearance, functionality, and homeostatic state. This homeostasis is dependent on an organized and timely renewal process, initiated by basal keratinocytes which proliferate and differentiate to ultimately transform into corneocytes that comprise the stratum corneum. An imbalance in this process has implications on skin’s appearance, health, and response to stress. Thus, it is essential to understand these changes to identify mechanistic intervention targets that would prevent and repair premature aging and maintain skin health and appearance.

We previously reported findings from a large base study that evaluated biopsies collected from photoexposed face and dorsal forearms as well as photoprotected buttock sites of Caucasian females across age decades spanning 20’s to the 70’s demonstrating that age impacts a wide range of molecular processes in skin.^2^ Given that a low grade chronic inflammatory state is hypothesized to be a significant contributor to premature aging in the inflammaging theory we asked whether this phenomenon could be observed in our previously reported skin biopsy study and investigated its potential impact on epidermal biology and homeostasis. A systems biology-based analysis of skin surface biomarkers, transcriptomics, proteomics, metabolomics, histology, and immunostaining confirmed that there is underlying chronic inflammation in photoexposed face skin that remains elevated across the decades. Primarily in photoexposed skin, we found there is an imbalance in epidermal homeostasis beginning in the 20’s to 30’s and elevation of senescence related components 30’s to 40’s. A subsequent increase of oxygen sensing/hypoxia and metabolic shift towards glycolysis occurs in the 50’s. Additionally, there is a higher epigenetic aging rate in 60’s when comparing to the 30’s and is further elevated by photoexposure. Based on these findings, we propose that photoexposed skin undergoes inflammaging which may play a role in the molecular and morphological changes that ultimately lead to a photoaged appearance and less healthy state of skin.

## 2 MATERIALS AND METHODS

The detailed protocols and statistical analysis are described in Supplemental Materials and Methods.

## 3 RESULTS

### 3.1 Age-associated changes in epidermal morphology

We first performed a histomorphometric analysis of the structural compartments of the epidermis from buttock, arm, and face sites across age groups. With age, the overall thickness of the stratum corneum increased (Figure 1A) whereas the epidermal layer becomes thinner (Figure 1B), and the rete ridge path length ratio decreases (Figure 1C). Comparison of the mean data between the 20’s and each decade showed that these changes become statistically significant in the older age groups (Supplemental Table 1). A representative histological stain from a 20’s and a 60’s year old face highlights these structural changes (Figure 1D). In an older age sample, we observed relatively lower detection of microcapillary structures using staining against UEA-1, a lectin that binds to endothelial related cells.^3^ A representative image shows the differential staining pattern below the basement membrane (Figure 1E, white arrows) as well as staining in the stratum granulosum and corneum. This pattern is similar to what has been previously reported in skin.^3^ The structural changes of thickening of the stratum corneum and the thinning of the epidermis suggest an imbalance between proliferation and differentiation that changes with age across all body sites.

**Figure 1.**
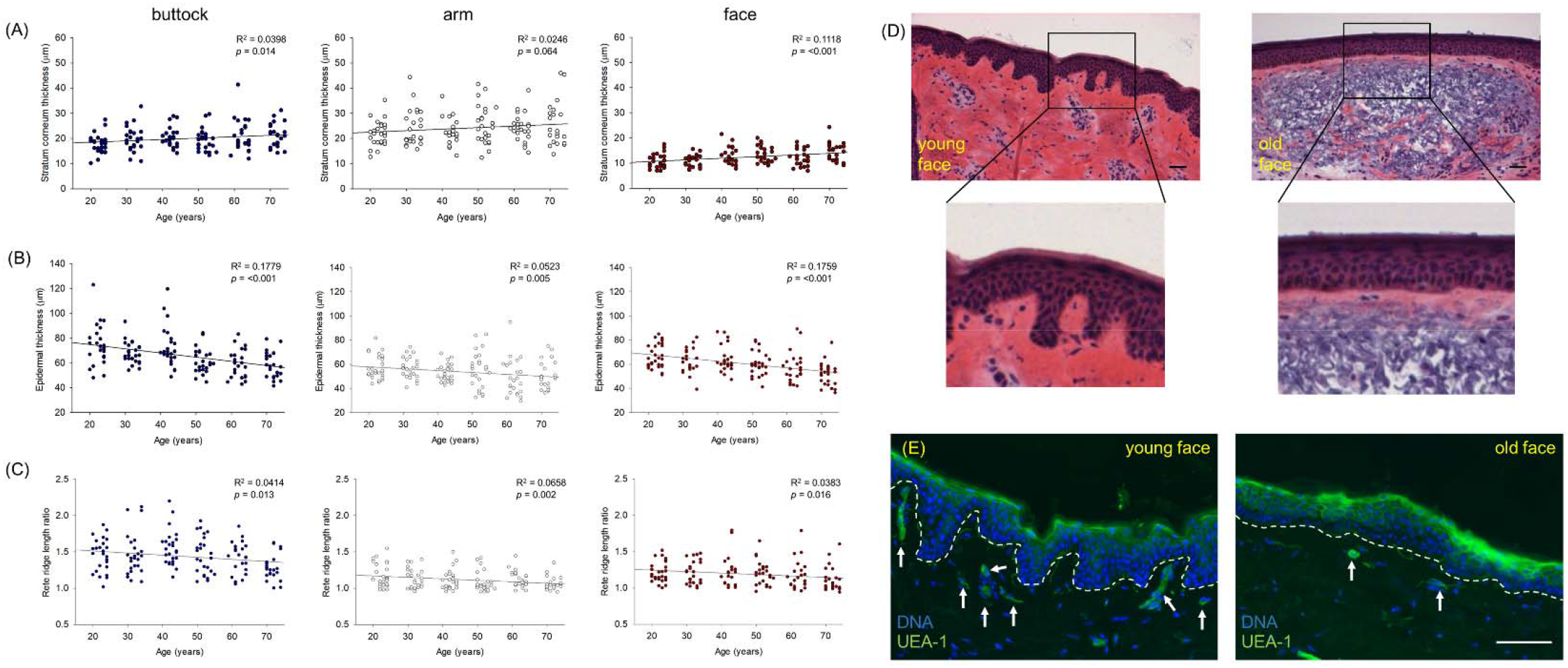
Age-associated changes in epidermal morphology. Epidermal structural elements were measured from hematoxylin and eosin stained histologic sections of skin from buttock, arm, and face body sites. Measurements are shown in scatter plots from all subjects and body sites for (A) stratum corneum thickness (μm), (B) epidermal thickness (μm), and (C) rete ridge length ratio (buttock – blue dots; arm – white dots; face – red dots). Linear regression analysis shows significant changes in structural morphology with age across all body sites. (D) Representative histological section from a 23-year-old and 72-year-old face site are shown along with the region of interest in black box as magnified panel to highlight stratum corneum thickening, epidermal thinning, and decrease of rete ridge length ratio in older aged face. Scale bar (black) is 100 μm. (E) Immunofluorescence staining for microcapillary beds (white arrows) by *Ulex Europaeus*-I Lectin (UVE-1, green) and DNA labelling (blue) show decreased staining of capillary bed structures in a representative section from a 23-year-old and 73-year-old face site. White dotted line denotes bottom of epidermal basal layer. Scale bar (white) is 100 μm.

### 3.2 Proteomics analysis shows elevated presence of proteins associated with differentiation and glycolysis in 60’s aged dorsal forearm epidermis over 20’s age group

To better understand these measured changes in epidermal structure with age, LCM isolated epidermal sections from 20’s and 60’s dorsal arms were processed and analysed by label free quantitative mass spectrometry. Out of 367 proteins identified, 83 showed a significant difference (*p*<0.1) in levels when comparing between the two age groups (Supplemental Table 2). Of the 83 proteins, 24 proteins were associated with epidermal differentiation and metabolism/oxygen sensing (Table 1). 23 of these had a similar directional relationship with their representative gene expression pattern with age. The exception is calpain 1 (CAPN1) which showed no significant change in expression levels across the decades (data not shown). Interestingly, we also detected a higher numerical level of hemoglobin-α (*p*=0.092) and hemoglobin-β (*p*=0.134, data not shown) present in the older group.

**Table 1.**
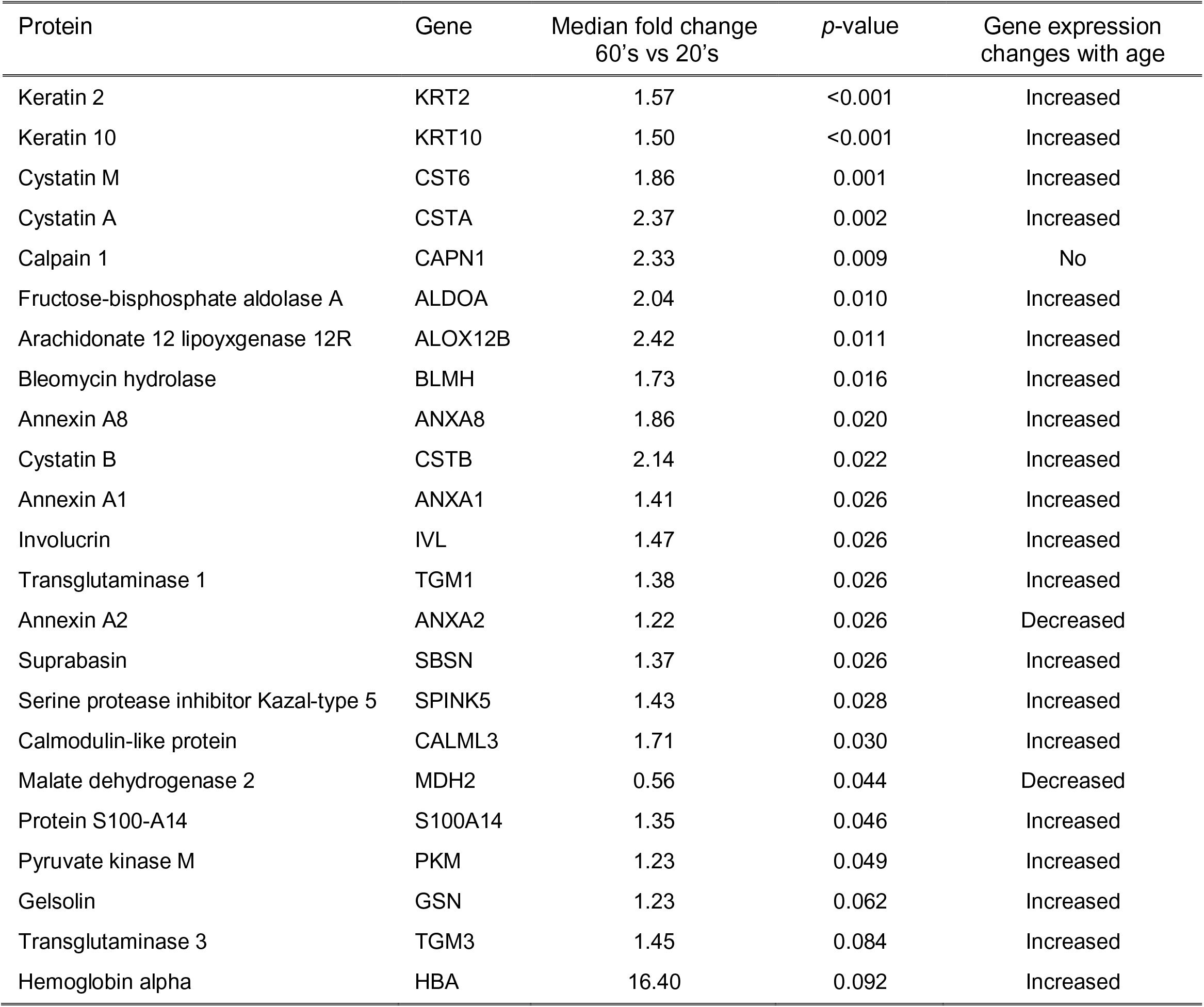
Median fold change of detected proteins between 60’s and 20’s age groups from laser capture microdissection sections of epidermis from photoexposed dorsal forearms and changes in gene expression correlation.

### 3.3 Imbalance in epidermal differentiation/proliferation increases with age in photoexposed epidermis

To further understand the age-associated changes in epidermal morphology and corresponding protein level changes, we manually curated transcriptomics data for genes encoding for proteins involved in epidermal differentiation and proliferation, including the epidermal differentiation complex, keratins, protease inhibitors, proteases, calcium binding proteins/AMP (antimicrobial peptides), proliferation, and late cornified envelope proteins (Figure 2A).^6, 7^ Statistical analysis of changes across the decades between 20’s and 70’s showed a pattern of elevated expression with age in the photoexposed dorsal arm and face sites in most of these groups (Figure 2A, pink coloration). In contrast, genes associated with proliferation showed a decline in expression with age decades across all three body sites (Figure 2A, blue coloration). Trace profiles of representative probe sets from face of filaggrin (FLG), involucrin (IVL), arachidonate 12-lipoxygenase,12R (ALOX12B), loricrin (LOR), keratin 2 (KRT2), keratin 14 (KRT14), calmodulin-like protein 3 (CALML3), serine protease inhibitor Kazai-type 5 (SPINK5), cystatin B (CSTB), Krüppel-like factor 9 (KLF9), insulin like growth factor 1 receptor (IGF1R), and late cornified envelope 2C (LCE2C) show the relative expression changes across the decades (Figure 2B). Interestingly, the late cornified envelope proteins did not show as significant of a pattern when comparing across 20’s and 70’s but exhibits a significant increase up to the 50’s and the reversal from 50’s to 70’s. To further visualize the gene expression profiles, we immunostained for several of these proteins in representative samples from young and old face and arm sites.

**Figure 2:**
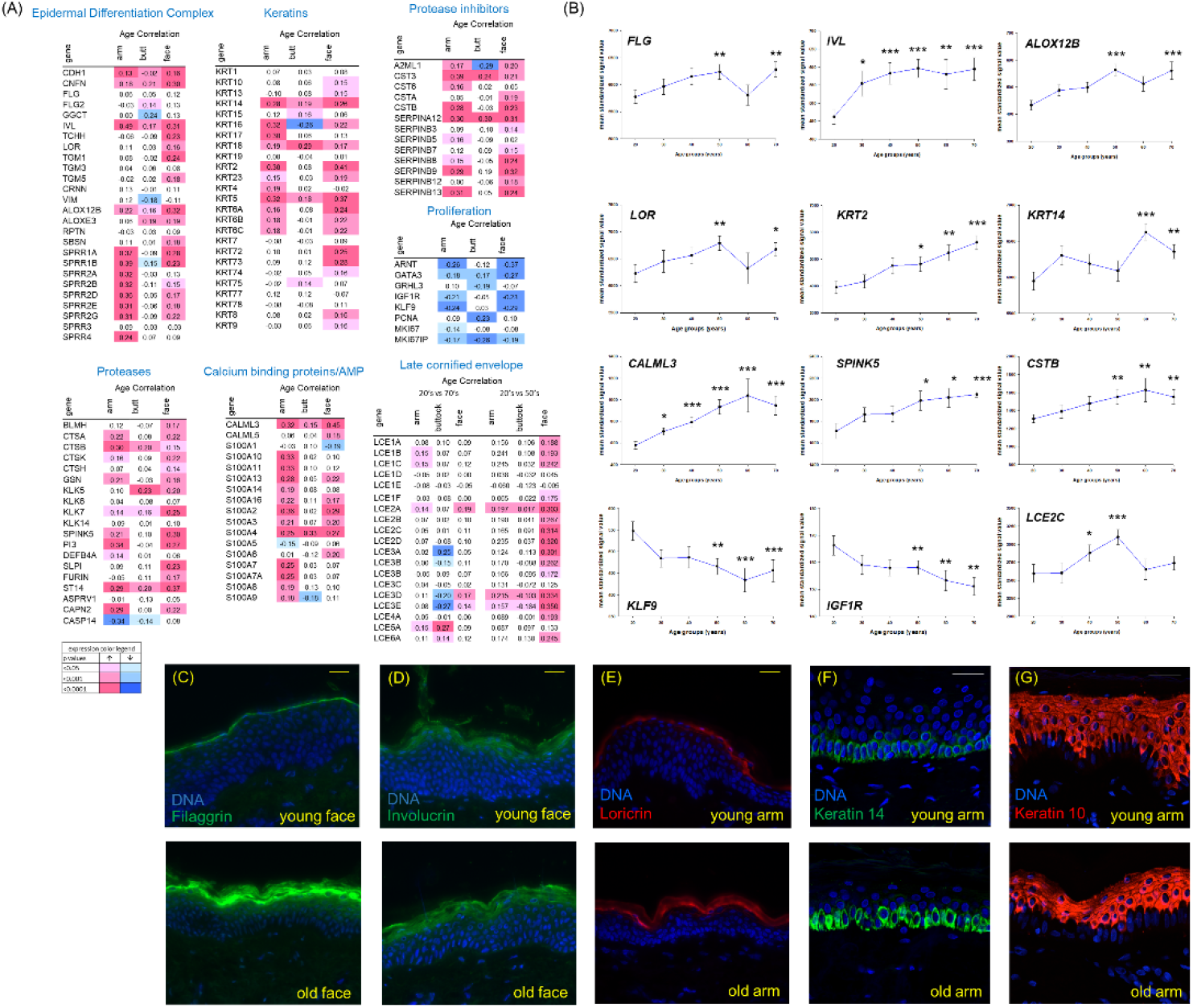
Transcriptomics profiling and immunofluorescent staining of epidermal differentiation genes and proteins with age. (A) Transcriptomics statistical heatmap of relative expression changes with age of select genes encoding for proteins associated with regulation of epidermal homeostasis. Genes associated with the epidermal differentiation complex, proteases, keratins, protease inhibitors, and calcium binding proteins/AMP (AMP - antimicrobial peptides) show a pattern of elevated expression across the decades from 20’s to 70’s (red). Genes regulating proliferation show a pattern of decreased expression across the decades from 20’s to 70’s (blue). Genes encoding for late cornified envelop proteins show a significant increase from the 20’s to 50’s (red) but was not significant between 20’s to 70’s. Color range reflects average normalized intensity values for each group (color legend panel: red higher expression and blue lower expression). (B) Trace profiles of probe sets from the face encoding for filaggrin (FLG), involucrin (IVL), arachidonate 12-lipoxygenase, 12R (ALOX12B), loricrin (LOR), keratin 2 (KRT2), keratin 14 (KRT14), calmodulin-like protein 3 (CALML3), serine protease inhibitor Kazai-type 5 (SPINK5), cystatin B (CSTB), Krüppel-like factor 9 (KLF9), insulin like growth factor 1 receptor (IGF1R), and late cornified envelope 2C (LCE2C). Significance indicates comparisons to 20-year-old cohort mean value (* *p*<0.05, ** *p*<0.01, *** *p*<0.001). Representative immunofluorescence images for comparison of filaggrin (C) and involucrin (D) from a 20-year-old (young face) and 64-year-old (old face) face site; loricrin (E), keratin 10 (F), and keratin 14 (G) from a 20-year-old (young face) and 60-year-old (old arm) arm site showing higher detection in older site. Yellow scale bar is 100 μm and white scale bar is 30 μm.

Immunostaining for filaggrin showed heightened levels in the upper granular/stratum corneum layers in a representative older age face site (Figure 2C) and to a lesser extent for involucrin and loricrin (Figure 2D and 2E). The basal keratin 14 marker showed a higher overall level of detection in a representative older age arm site (Figure 2F) and a modestly higher level of detection of the suprabasal marker keratin 10 (Figure 2G).

### 3.4 The IL-1RA/IL-1**α** ratio and IL-8 remain elevated across the decades in photoexposed facial skin, the *cis/trans* urocanic acid and 53BP1 DNA damage foci are detected in photoexposed sites, and epigenetic age is higher with age in photoexposed arm sites

In addition to proteomics analysis on LCM derived epidermal sections we tested for the presence of the senescence-associated secretory phenotype (SASP) inflammatory biomarkers IL-8 and the IL-1RA/IL-1α ratio on the surface of skin.^8^ Analysis of tape strip extractions showed the levels of both biomarkers were elevated in photoexposed face compared to dorsal arm and buttock sites (Figure 3A and 3B). Interestingly, the levels on face remained elevated across the age groups. It was surprising that we did not detect elevated levels of these cytokines in the photoexposed dorsal arm sites. To better understand this difference between the two sites, we analysed for the UV-sensitive metabolite ratio of *cis/trans*-urocanic acid. We showed a significant elevation in both face and arm compared to buttock site and was consistent across the decades (Figure 3C). We also stained arm and buttock sites from both young and old for 53BP1, an indicator of DNA damage response induced by UV-irradiation.^9, 10^ Quantification showed significantly more foci in the basal layer of aged arm compared to young, while very few foci were detected in buttock (Figure 3D-F). The buttock sites in either young or old did not show any significant increase in DNA damage. Additionally, we quantitated epigenetic age levels, which is based on DNA methylation levels from thousands of aging related locis.^11^ We showed that both body sites showed elevated epigenetic age levels in the 60’s when compared with the 30’s and significantly accelerated aging in arm sites compared to buttock sites (Figure 3G). The elevated levels of these photosensitive markers with age in arm and face sites support that both sites undergo a certain degree of photodamage. The muted levels of IL-8 and the IL-1RA/IL-1α ratio may be due to unknown physiological differences that merit further investigation.

**Figure 3:**
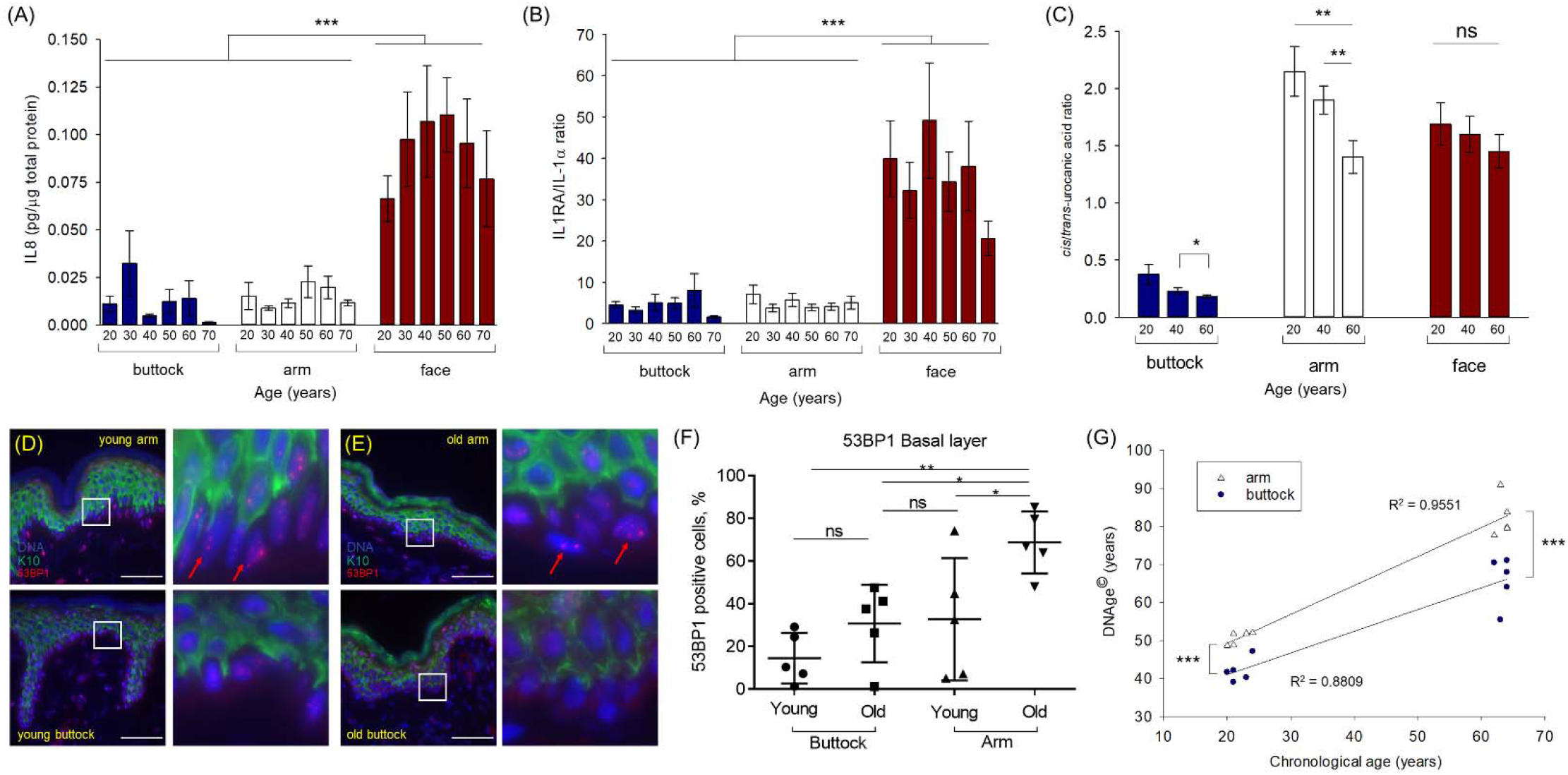
Markers of inflammation, photoexposure, and DNA damage and methylation comparisons across body sites and age. Biological material extracted from tape strips collected across all sites and age groups were analyzed. (A) IL-1RA/IL-1α and (B) IL-8 are reported from dorsal forearm, cheek, and buttock sites across each decade age group showing elevated levels present in face sites across all decades. (C) The *cis/trans*-urocanic acid ratio are reported from dorsal forearm, cheek, and buttock sites from 20’s, 40’s, and 60’s age groups and show elevated levels across decades in both arm and face sites. Representative immunofluorescence images for detection of foci localization of the DNA damage marker 53BP1 (red) in a 20-year-old young arm and buttock (D) and a 64-year-old arm and buttock (E) are shown. Keratin 10 staining (green) and DNA labelling (blue) are shown. Region of interest outlined in white is shown as magnified panel and red arrows indicate examples of elevated 53BP1 foci in nuclei of older photoexposed site. Scale bar, 100 μm. (F) Quantification of 53BP1 positive cells in K10-negative basal layer from young and old arm and buttock images (n=5 per site). Data is shown as mean ± SD. Ordinary one-way ANOVA was performed with Sidak’s multiple comparison test. (G) Quantitation of epigenetic age levels based on DNA methylation levels from young and old arm and buttock images (n=5 per site). * *p*<0.05, ** *p*<0.01, *** *p*<0.001, ns non-significant.

### 3.5 Senescence and inflammation are elevated with age in photoexposed epidermis

The detection from facial skin surface of elevated levels of IL-8 which remains consistently high (Figure 3B) suggests that this site may present a higher senescence/inflammation rate than arm. As shown previously, we report that the senescence associated gene CDKN2A was elevated with age across all three body sites.^2^ To better understand the correlation there may be between senescence, the heightened presence of the SASP-associated inflammatory biomarkers, and the epidermal morphological changes, we manually curated transcriptomics data for a subset of genes encoding for senescence and inflammatory associated proteins. We found an overall pattern of increased expression across the decades between 20’s and 70’s in both the photoexposed arm and face sites, with more genes being upregulated in face (Figure 4A). In agreement, cyclin dependent kinase inhibitor 2A (CDKN2A), alpha-crystallin B chain (CRYAB), cytokine receptor type 2/IL8RB (CXCR2) were upregulated upon aging (Figure 4B). Similarly, several genes that have been reported to be reduced upon senescence (RBL2, SIRT, LMNB1) showed a general pattern of lowered expression (Figure 4A).^12–14^ CDKN2A is known to encode for several proteins involved in senescence and linkages to cancer, and aging, including p16^INK4A^.^15–17^ To further visualize the expression patterns, we immunostained biopsy sections from young and old face sites for p16^INK4a^ and observed higher number of p16-positive cells in age facial skin (Figure 4C and 4D, yellow arrows).

**Figure 4:**
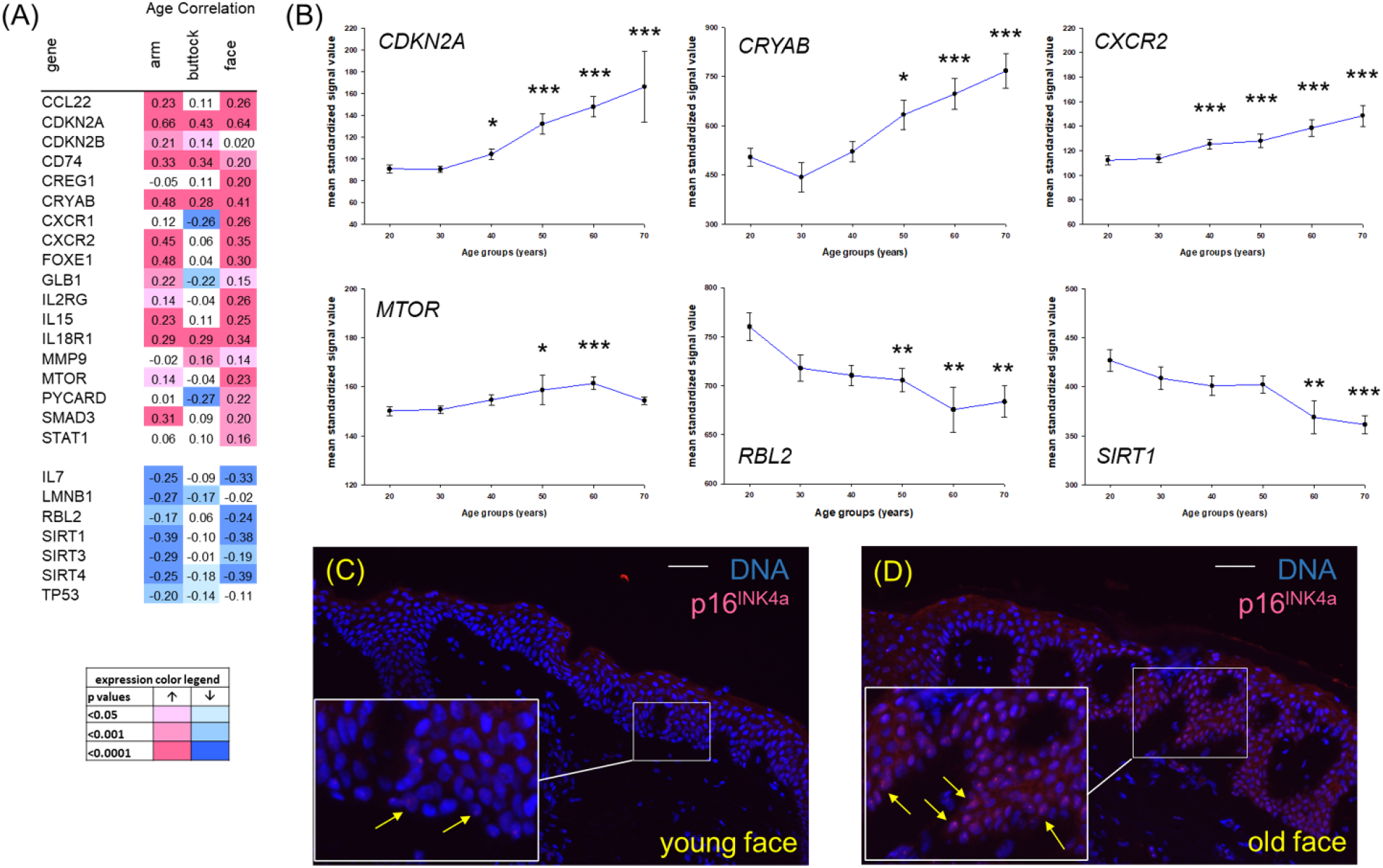
Transcriptomics profiling of senescence associated genes and immunofluorescent staining for p16^INK4a^. (A) Transcriptomics statistical heatmap of relative expression changes with age of select genes encoding for proteins associated with regulation of senescence. Genes associated with increasing cellular senescence and senescence associated secretory phenotype show a pattern of elevated expression across the decades from 20’s to 70’s in the photoexposed sites of dorsal forearm and face (red). Additionally, genes associated with mitigating senescence show a pattern of decreased expression across the decades from 20’s to 70’s (blue). Color range reflects average normalized intensity values for each group (color legend panel: red higher expression and blue lower expression). (B) Trace profiles of probe sets from the face encoding for cyclin dependent kinase inhibitor 2A (CDKN2A), alpha-crystallin B chain (CRYAB), cytokine receptor type 2/IL8RB (CXCR2), mammalian target of rapamycin (MTOR), retinoblastoma-like protein 2 (RBL2), and sirtuin 1 (SIRT1). Significance indicates comparisons to 20-year-old cohort (* *p*<0.05, ** *p*<0.01, *** *p*<0.001). Representative immunofluorescence images for staining of p16^INK4a^ (pink) and DNA labelling (blue) from a 21-year-old (C, young face) and a 63-year-old (D, old face) face site. Region of interest outlined in white is shown as magnified panel and yellow arrows indicate examples of elevated p16 detection in nuclei of from older face site. Scale bar (white) is 100 μm.

### 3.6 An oxygen sensing/hypoxic fingerprint and metabolic reprograming increases with age in epidermis

The proteomics-based detection of elevated levels of several glycolytic enzymes in 60’s aged epidermal arm LCM samples suggests the epidermis was undergoing a metabolic shift. A shift to glycolysis is a hallmark process of cells when exposed to hypoxic conditions. The morphological changes with age of increased stratum corneum thickness, decrease in rete ridge path length ratio, and less vasculature detection could impact oxygen bioavailability in the epidermis. Finally, the increased expression and protein detection of hemoglobin-α further suggests an oxygen sensing response by the epidermis. Thus, we manually curated from transcriptomics data a subset of genes encoding for proteins sensitive to oxygen tension or associated with cellular responses to hypoxia. These genes showed an increased expression pattern across the decades between 20’s and 70’s in arm, buttock, and face sites (Figure 5A, pink coloration).

**Figure 5:**
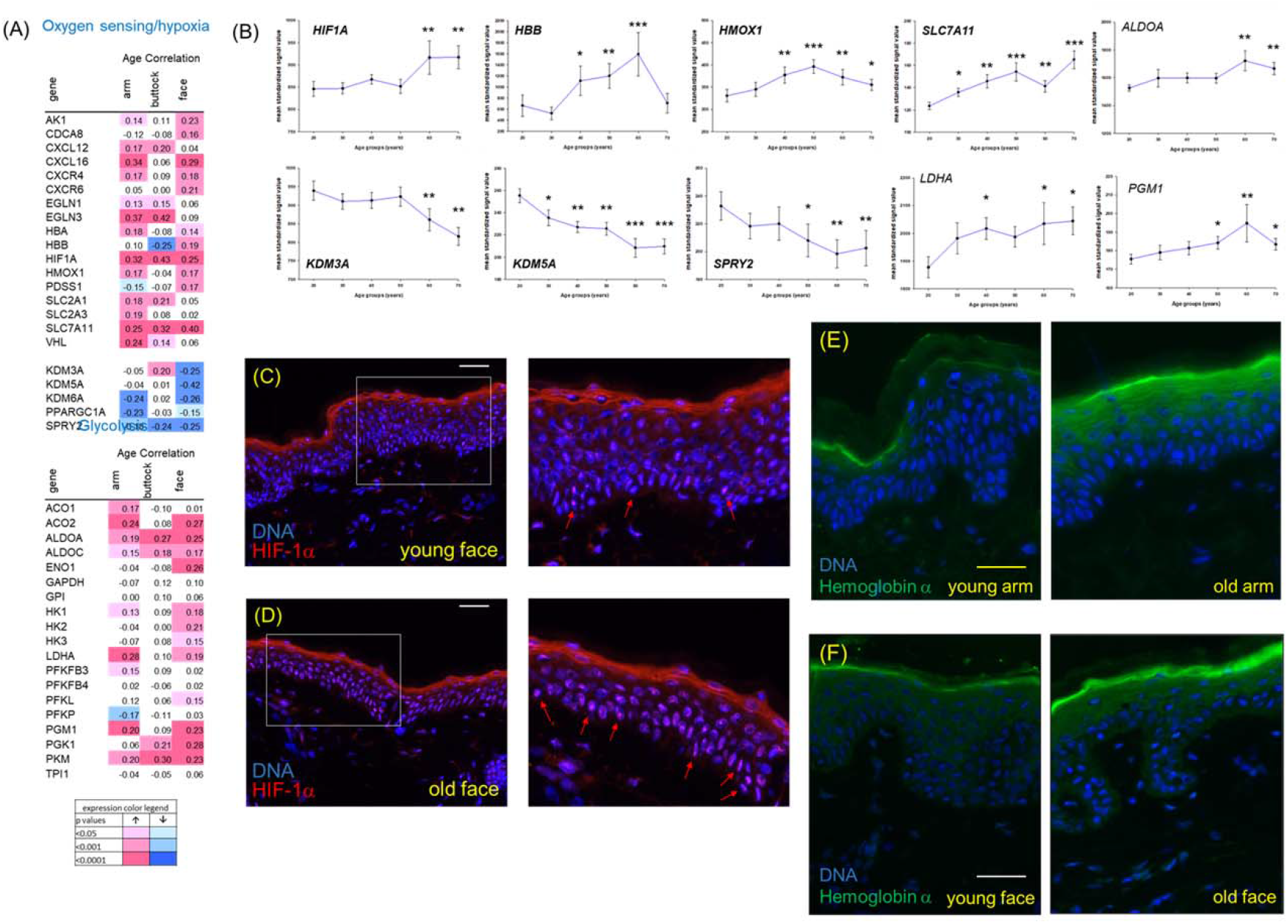
Transcriptomics profiling of oxygen sensing and hypoxia associated genes and immunofluorescent staining for HIF-1α and hemoglobin α. (A) Transcriptomics statistical heatmap of relative expression changes with age of select genes encoding for proteins associated with oxygen sensing and hypoxia. These genes show a pattern of elevated expression across the decades from 20’s to 70’s in the photoexposed sites of dorsal forearm and face (red). Additionally, genes associated with being downregulated under hypoxic conditions show a pattern of decreased expression across the decades from 20’s to 70’s (blue). Color range reflects average normalized intensity values for each group (color legend panel: red higher expression and blue lower expression). (B) Trace profiles of probe sets from the face encoding for hypoxia inducible factor 1, subunit alpha (HIF1A), hemoglobin-β (HBB), heme oxygenase 1 (HMOX1), cystine/glutamate antiporter (SLC7A11), aldolase A (ALDOA), lysine demethylase 3A (KDM3A), lysine demethylase 5A (KDM5A), Sprouty homolog 2 (SPRY2), lactate dehydrogenase A (LDHA), and phosphoglucomutase 1 (PGM1). Significance indicates comparisons to 20-year-old cohort (* *p*<0.05, ** *p*<0.01, *** *p*<0.001). Representative immunofluorescence images for comparison of HIF-1α between a 22 year old (C, young face) and a 60-year-old (D, old face) face sites. Region of interest outlined in white is shown as magnified panel and red arrows indicate examples of elevated HIF-1α staining in nuclei of older site. Scale bar (white) is 100 μm. Representative immunofluorescence images for comparison of hemoglobin-α between a 22-year-old (E, young arm) and a 63-year-old (E, old arm) arm sites and between a 21 year old (F, young face) and a 63 year old (F, old face) face sites showing elevated detection throughout the granular layer stratum corneum in the older sites. White scale bar is 25 μm and yellow scale bar is 50 μm.

Consistent with this, genes encoding for proteins known to negatively respond to hypoxia showed decreased expression across the decades, primarily in the photoexposed forearm and face sites (Figure 5A, blue coloration). Genes encoding for glycolytic enzymes were also analysed and several genes were found to have elevated expression patterns across the decades between 20’s and 70’s in arm and face sites (Figure 5A, red coloration). To further illustrate the statistical findings, representative expression traces are shown for hypoxia inducible factor 1, subunit alpha (HIF1A, a master regulator of cellular response to hypoxic conditions), hemoglobin-β (HBB), heme oxygenase 1 (HMOX1), cystine/glutamate antiporter (SLC7A11), aldolase A (ALDOA), lysine demethylase 3A (KDM3A), lysine demethylase 5A (KDM5A), Sprouty homolog 2 (SPRY2), lactate dehydrogenase A (LDHA), and phosphoglucomutase 1 (PGM1) (Figure 5B). To further understand HIF1A and hemoglobin gene expression, we immunostained for HIF-1α and hemoglobin-α. A representative image shows staining of HIF-1α in nuclei of young face sites but higher expression was detected in older aged face sites (Figure 5C and 5D, red arrows). Representative images of hemoglobin-α staining in both arm and face sites highlight an elevated staining intensity throughout the upper granular/stratum corneum layers in arm (Figure 5E) and face (Figure 5F) from older individuals as compared to younger. Interestingly, there was no observable staining increase in the basal layer and through the dermis, further supporting that the presence of hemoglobin-α was epidermally derived and not erythroid.

## 4 DISCUSSION

The skin is the first line of defense protecting the body from environmental stressors such as solar radiation and pollution. Daily exposure to sunlight is one of the more significant environmental insults that induces DNA damage, oxidative stress, and inflammation in skin. Human skin must maintain robust repair capabilities to prevent cumulative damage triggered by these stressors. However, with age this ability is diminished, and the onset of senescence further hinders the skin’s capacity to mitigate stress-induced inflammation and can lead to the presence of chronic low-level inflammation.^18^ This phenotype is a key feature of inflammaging. The evidence for the presence of inflammaging in skin has been previously reviewed and it was highlighted that while there are clear signs of an inflammaging microenvironment in skin, further work is needed to better understand it’s role on skin aging.^19^

To better understand the role inflammaging on skin aging, we utilized a systems-biology based approach to investigate biological samples collected from photoprotected and exposed female body sites spanning 6 decades of age. A previous report found that patterns of gene expression accelerated with aging in Caucasian females and differed in a subgroup that appeared exceptionally youthful based on image analysis of facial appearance.^2^ The current study focused on the epidermal skin compartment and employed a systems biology-based approach to increase our understanding and identify potential intervention strategies to mitigate premature aging. Our findings provide a body of evidence that photoexposed facial skin is in an inflammaging microenvironment due to the presence of elevated chronic inflammation which, in turn, could be a factor that leads to an imbalance in epidermal homeostasis starting in the 30’s as measured via histology, transcriptomics, and proteomics (Figure 6). This suggests that targeting inflammation in younger aged skin may be a promising intervention approach to mitigate the molecular and morphological changes that lead to a photoaged appearance of skin and impact on underlying skin health.

**Figure 6:**
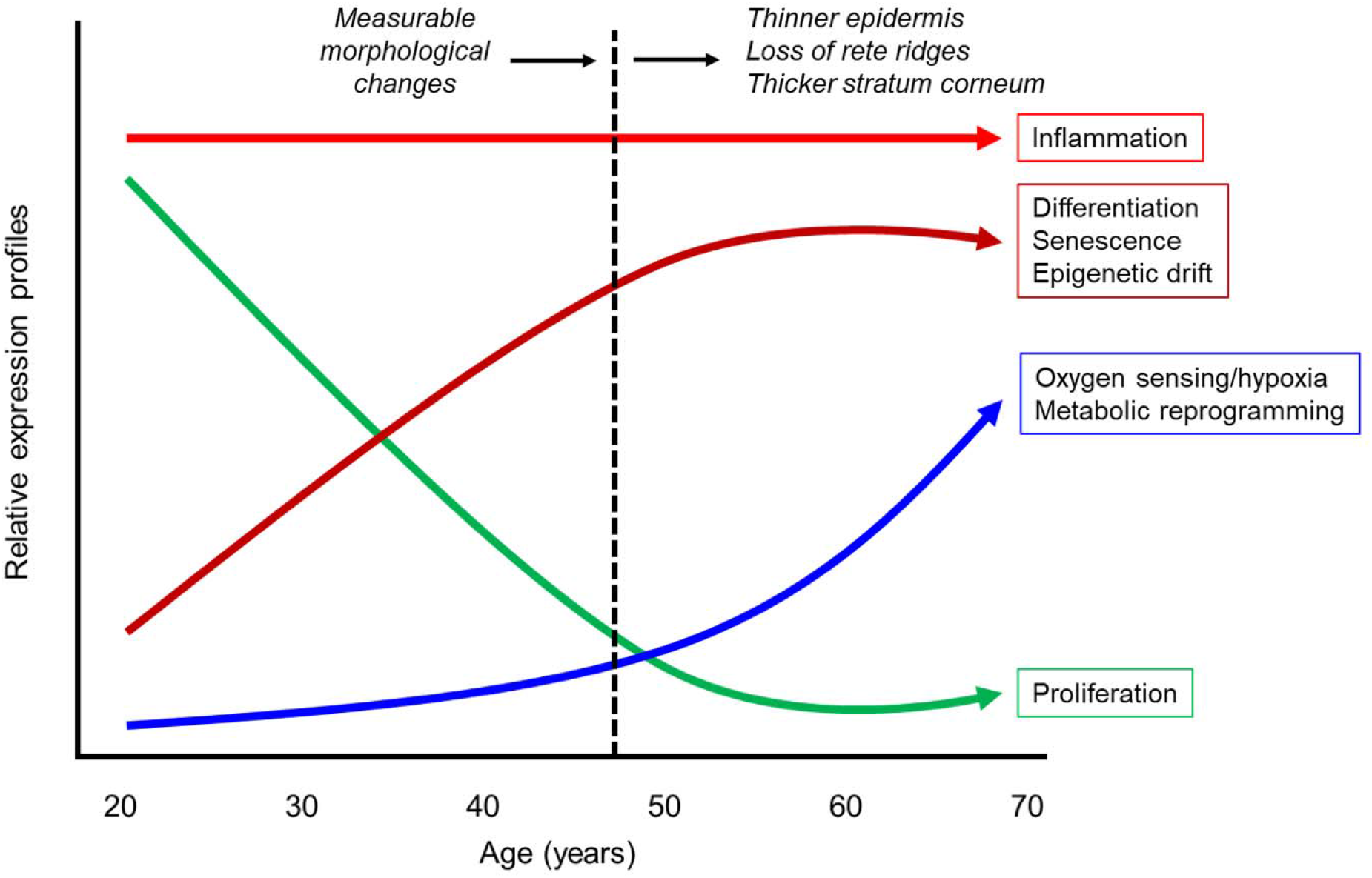
Schematic representation of key biological processes changing with age in epidermis of photoexposed facial skin. A heightened inflamed status remains constant across the decades in the epidermis of photoexposed facial skin and underlying processes such as differentiation, senescence, oxygen sensing/hypoxia, epigenetic drift, and metabolic reprogramming steadily increase. In contrast, epidermal proliferation declines with age. These suggest that inflammaging leads to an imbalance in epidermal homeostasis with photoaging.

The histomorphologic analysis in this study found that the epidermis undergoes significant changes with age, including stratum corneum thickening, implying that there may be a stronger barrier in older aged skin. While counter-intuitive, several reported studies have shown that trans-epidermal water loss values decrease in older aged subjects, suggesting that the barrier integrity improves with age.^20^ However, the underlying health of the skin plays a role to ensure optimal repair response kinetics to damaging agents. Older aged skin has been shown to have a slower and weaker response profile to damage such as wounding and tape strip removal.^21, 22^ We also show that with age the epidermis becomes thinner, the rete ridge path length flattens, and these changes correlate with changes in gene expression and protein levels associated with differentiation and proliferation. Expression changes occur in a large proportion of genes encoding for proteins associated with the epidermal complex, keratins, proteases, protease inhibitors, calcium bindings proteins/AMP, and late cornified envelope proteins. Additionally, these changes are more apparent in the photoexposed arm and face sites than the buttock site. This imbalance in differentiation and proliferation processes appears to shift in the 30’s and could be a factor in the observed morphological changes detected starting in the 40’s. For example, the representative expression traces for FLG, LOR, ALOX12B, KRT2, CALML3, SPINK5, and CSTB all show a similar pattern of increased expression beginning in the 20’s to 30’s and continuing to increase across the decades. It is worth noting that some of these markers show alterations of this trend in the 50’s, presumably due in part to hormonal changes as recorded in the previous study.^2^ This is particularly highlighted in the respective traces presented as well as the overall expression patterns for the late cornified envelope proteins which showed significant changes in expression between the 20’s and 50’s but lost significance when comparing between the 20’s and 70’s.

Several of the proteins expressed by these genes were also detected via proteomics profiling between the photoexposed arm of young and old subjects. A similar proteomics profiling has been reported in which the authors used tape strip collection to quantitate the levels of surface proteins associated with differentation.^23^ Their findings are similar to the ones presented here with the exception that several proteins showed contrasting reduced levels in photoexposed skin compared to the elevated levels of those same proteins in our study. Overall, there is an apparent correlation between the differentiation associated gene expression changes that begins in the 20’s and correlates with the morphological changes that become significantly measurable starting in the 40’s. This suggests an imbalance in epidermal homeostasis which could impact its response profile to environmental insults and maintenance of normal cellular function.

To better understand the inflammatory and photoexposure status of the subjects in this study, we evaluated for the presence of inflammatory and photosensitive biomarkers isolated from the skin’s surface. Detection of elevated levels of IL-8 has been shown to be elevated in eczema, atopic dermatitis, and psoriasis skin and in 3D skin models after UVB exposure ^24–27^ We found elevated levels of IL-8 on photoexposed facial skin surface sites that remain elevated across age groups. The ratio of IL-1RA/IL-1α present on the skin’s surface is known to be an indicator of underlying inflammation associated with skin dermatitis conditions and UV exposure.^28–30^ Relative to impact of age and photoexposure on this inflammatory biomarker, it was reported that the IL-1RA/IL-1α ratio was elevated in photoexposed face compared to non-exposed upper inner arm and remained constant across age groups.^28-30^ Relatedly, we show similar patterns when comparing between photoexposed face where the IL-RA/IL-lα ratio was consistently high and consistently low in photoprotected buttocks across the decades. Surprisingly, we did not see an increase in these cytokines in photoexposed dorsal arm samples since we had previously reported there are significant histological indications of photoaging.^2^ We show that several biomarkers associated with photoexposure are increased in arm sites, including the *cis/trans*-urocanic acid ratio, foci of the DNA damage response marker 53BP1 that is sensitive to UV exposure, and epigenetic age derived from methylation levels of DNA an indicator of epigenetic aging.^11, 31–33^ These methylation patterns are similar to what has been previously reported where the biopsies were enzymatically separated into epidermis and dermis fractions in contrast to LCM in our study.^34^ Overall, this supports that the photoexposed arms undergo photodamage. We do not believe the lower levels of IL-8 or the IL-1RA/IL-1α ratio on photoexposed arm or buttock sites are an artefact since we performed the analysis in two independent experiments from duplicate tapes. The difference could reflect a dose response or a level of chronic exposure or, alternatively, facial skin is among the thinnest in the body and may be more susceptible to injury. While overall our results support the hypothesis that photoexposed skin is in a heightened state of inflammation, and that inflammation is present early in the 20’s and remains persistent across the decades, future work is needed to understand the physiological relevance in photodamaged arms. Overall, the implications of this constant inflammatory pressure could be an indicator of skin inflammaging that leads to the changes in gene expression patterns and correlating protein levels.

We previously reported CDKN2A, a gene that encodes for proteins associated with senescence induction, to be elevated with age.^2^ CDKN2A is known to encode for p14^ARF^, p15^INK4B^, and p16^INK4A^, all of which are involved in senescence and play significant roles in cancer, and aging, including in skin.^15–17^ In the current study we wished to better understand this correlation beyond CDKN2A and performed a focused transcriptomics profiling of select genes encoding for proteins associated with regulation or induction of senescence in skin.^35^ It has been established that photoexposure can cause keratinocytes to prematurely enter senescence and these cells can be characterized by secretion of an altered secretome called the senescence-associated secretory phenotype (SASP) and is enriched with pro-inflammatory cytokines such as IL-6, IL-8, and IL-1β.^8^,

The elevated skin surface levels of IL-8 early in the 20’s age cohort on photoexposed face sites supports there may be an early onset of a SASP associated phenotype in photodamaged facial skin. We see significant elevated levels of expression of genes encoding for proteins associated with senescence in the photoexposed sites. For example, GLB1 encodes for SA-β-gal (beta-galactosidase), a well-known biomarker of senescence in numerous tissues, including skin.^35^ Several chemokine receptors were observed to increase in expression levels with age in the photoexposed arm and face sites. CXCR1 and CXCR2 encode for receptor proteins that bind with IL-8 and showed elevated expression in both arm and face.^36^ Interestingly this provides a potential correlation of inflammatory response with the elevated levels of IL-8 present on the skin’s surface. A survey of candidate SASP components from a comparison between *in vitro* senescence models and *in vivo* tissue and fluid samples showed the elevated presence of CCL22, IL15, and MMP9 under senescent-impacted conditions.^37^ The mammalian target of rapamycin (mTOR) is suggested to be a master regulator of metabolite sensing that impacts senescence induction and overall cellular aging.^38, 39^ We show in both photoexposed epidermal sites an increase in mTOR expression levels with age (Figure 4A) that becomes significant in the 50’s compared to the 20’s for face (Figure 4B). CREG (cellular repressor of E1A-stimulated genes 1) co-expression with p16^INK4a^ can further enhance senescence than either expressed alone.^40^ Recently, CRYAB and HMOX1 have been proposed to be senolytic targets in humans cell models.^41^ Interestingly, it was also demonstrated that HMOX1 expression levels were increased during differentiation, which supports a similar correlation as measured in our study.^42^ As reported here, we observe a significant increase in the expression patterns of these genes starting in the 30’s and continuing into the 70’s in photoexposed facial epidermis sites. In addition, we evaluated genes that encode for proteins that mitigate senescence, including RBL2, SIRT1, SIRT3, SIR4, and TP53.^12, 43–45^ These show varying patterns of decreased expression in the epidermis of photoexposed sites with significance starting in the 50’s and 60’s. Finally, we immunostained for p16^INK4A^ and detected nuclear localized puncti in both basal and spinousal layers. In total, the data presented here supports that photoexposed skin is undergoing an accumulation of senescent cells with age. The chronic presence of the SASP factor IL-8 could be a causative indicator of senescence but further work is needed to establish cause and effect linked to the imbalance in differentiation-/proliferation and morphological changes.^46^ The implications of skin undergoing these changes in inflammation and senescence due to photoexposure also has potential implications on overall body health. A recent review suggests there is a correlation between the accumulation of senescent cells in the skin and a negative impact on overall systemic health and longevity that occurs via the hypothalamic-pituitary-adrenal axis.^47^

Oxygenation of the epidermis occurs via passive diffusion from direct contact with atmospheric oxygen and from microcapillary beds intertwined underneath the basement membrane.^48^ This may explain why the epidermis is considered to have a relatively low oxygen tension that has been estimated to range between 0.3-8% and why the epidermis could be considered hypoxic in contrast to the highly vascularized dermis where oxygen levels are estimated to be >7%.^49, 50^ The morphological changes that were measured in the epidermis with age suggested to us there could be a further limitation of oxygen supply due to the longer diffusion path length through the thickened stratum corneum as well as the reduced surface area interface with microcapillary beds from reduction of rete ridge undulation pattern. It has been previously reported that aging can lead to a measured increase in hypoxic-related response profiles.^51^ That work utilized suction fluid blisters from young and older aged upper arms for transcriptomics profiling. In our study, we utilized the sensitivity of LCM dissection to localize the epidermis in both photoprotected and photoexposed skin sites for further investigation and an overall systems-biology body of evidence. In addition to HIF1A, an expanded transcriptomics profiling of select genes encoding for proteins associated with regulation or responsiveness to oxygen tension changes or hypoxia supports our hypothesis that photoaged skin is transitioning into a more hypoxic microenvironment. For example, hypoxic conditions have been shown to induce HMOX1 gene expression at 1% O_2_ *in vitro* and 7% O_2_ *in vivo* and this was mediated by HIF-1α activity.^52^ Gene expression of the CXCL16-CXCR6 axis, CXCR4, and CXCL12 have been reported to be elevated under chronic hypoxic conditions.^53^ PDSS1 encodes for decaprenyl diphosphate synthase subunit 1 and was recently identified as a member of a hypoxia signature in hepatocellular carcinoma cells.^54^ We identified several genes whose expression patterns are negatively regulated under hypoxic conditions. Lysine demethylase 3A (KDM3A) has been reported to regulate PGC1α (PPARGC1A) and is inhibited under hypoxic conditions.^55^ Silencing of SPRY2 gene expression was shown to correlate with elevated levels of HIF-1α.^56^ Prolonged exposure to hypoxic conditions is known to shift cellular metabolism to a greater reliance on glycolysis due to the more anaerobic conditions.^57^ We observed a similar shift based on elevated expression of genes encoding for enzymes involved in glycolysis such as ALDOA, ENO1, LDHA, PGM1, and PKM. This was further supported by the detection of higher protein levels for ADLOA and PKM in older aged arm samples compared to younger aged samples. Expression for the glucose transporters SLC2A1, SLC2A3, and SLC7A11 were also elevated with age, which have been reported to be stimulated in response to hypoxia.^54, 58^ Interestingly, we detected elevated expression of hemoglobin-α and -β (HBA and HBB) and a numerically greater level of hemoglobin-α protein levels in older aged photoexposed arms. Of note, we did not see any significant staining for hemoglobin-α through the dermis and neither did we identify hemoglobin differences from proteomics of dermal sections (data not shown). While hemoglobin is well known for its role in O_2_ and CO_2_ gas exchange in red blood cells, an increasing number of non-erythroid tissues have been reported to endogenously express hemoglobin.^59^ The exact function of hemoglobin in non-erythroid tissue is not clear but it has been speculated it could include regulation of heme, iron and oxygen levels.^60^ It has also been proposed that hemoglobin plays a role in response to oxidative stress by helping protect against ROS damage.^61^ Overall, the significant increase in expression of genes associated with hypoxia and glycolytic enzymes suggests a phenotype reflecting a hypoxic microenvironment in photoexposed skin and, to a weaker extent, in non-exposed skin. Further work is needed to validate these findings with quantitation of differences in oxygen content in the epidermal compartment as a function of age and photoexposure.

Limitations exist in this study since it is not clear on the causal relationship between the molecular changes ascribed and the cascade across the decades to the morphological changes.

## 5 CONCLUSIONS

In summary, this systems biology-based approach to analyse inflammatory and photosensitive biomarkers, proteomics, transcriptomics, and immunostaining strongly suggests that photoexposed facial skin is undergoing inflammaging that begins as early as in the 20’s and that multiple biologic pathways are affected in this process. We propose that the chronic presence of inflammation and SASP early in age may contribute to the molecular reprogramming, imbalance of epidermal homeostasis, and morphological changes. The presence of heightened senescence, oxygen sensing/hypoxic response, epigenetic drift, and metabolic shift may also play roles leading to this imbalance. While this work provides further evidence on the role of senescence in impacting skin aging, further evidence is still required.^62, 63^ Finally, the detection of non-erythroid-derived hemoglobin in the epidermis is a novel finding that merits further evaluation on its function and role in skin biology and aging.

## Abbreviation list

LCM: laser capture microdissection
SASP: senescence-associated secretory phenotype
DEJ: dermal epidermal junction
UEA-1: *Ulex Europaeus*-I Lectin
53BP1: p53-binding protein 1
IL-8: interleukin-8
IL-1α: interleukin-1α
IL-1RA: interleukin-1 receptor antagonist
FLG: filaggrin
INV: involucrin
ALOX12B: arachidonate 12-lipoxygenase,12R
LOR: loricrin
KRT2: keratin 2
KRT14: keratin 14
CALML3: calmodulin-like protein 3
SPINK5: serine protease inhibitor Kazai-type 5
CSTB: cystatin B
KLF9: Krüppel-like factor 9
IGF1R: insulin like growth factor 1 receptor
LCE2C: late cornified envelope 2C
CAPN1: calpain 1
CDKN2A: cyclin dependent kinase inhibitor 2A
CRYAB: alpha-crystallin B chain
CXCR2: cytokine receptor type 2/IL8RB
mTOR: mammalian target of rapamycin
RBL2: retinoblastoma-like protein 2
SIRT1: sirtuin 1
HIF1α: hypoxia inducible factor 1, subunit alpha
HBA: hemoglobin-α
HBB: hemoglobin-β
HMOX1: heme oxygenase 1
SLC7A11: cystine/glutamate antiporter
ALDOA: aldolase A
KDM3A: lysine demethylase 3A
KDM5A: lysine demethylase 5A
SPRY2: Sprouty homolog 2
LDHA: lactate dehydrogenase A
PGM1: phosphoglucomutase 1

## ACKNOWLEDGEMENTS

We acknowledge the assistance of John C. Bierman in facilitating tape strip analysis.

## AUTHOR CONTRIBUTIONS

BBJ, CYRT, CYH, TTL, XY, LC, SP, SB, OD, and JEO conceived the experiments; BBJ, CYRT, CYH, ALS, TTL, XY, CN, WG, YCC, YMD, LC, and PSG performed experiments and analysed the data. JEO wrote the manuscript. All authors reviewed/edited the manuscript.

## DATA AVAILABILITY STATEMENT

The data that support the findings of this study are available from the corresponding author upon reasonable request.

