## Supplementary material for "Inflammaging in human photoexposed skin: Early onset of senescence and imbalanced epidermal homeostasis across the decades": Methods and Materials

**Supplementary Materials and Methods**

**S.1 Human clinical sample collection and data availably**

Collection of skin biopsies and processing for mRNA extraction, target labeling, processing, and analysis has been previously described.^2^  The data reported in this paper have been deposited in the Gene Expression Omnibus (GEO) database, <https://www.ncbi.nlm.nih.gov/geo> (accession no. [GSE112660](https://www.ncbi.nlm.nih.gov/geo/query/acc.cgi?acc=GSE112660)). No other transcriptomics datasets were generated or analyzed for this manuscript.

**S.2 Histology and histomorphometry**

Histomorphometry was conducted to observe age-associated skin structural alterations.  Hematoxylin and eosin staining was performed on 10 µm cryosections from fresh frozen skin biopsies with the Shandon Rapid-Chrome Frozen Section Staining Kit (Thermofisher Scientific, Kalamazoo, MI) according to manufacturer’s recommendations.  Sections were fixed with Rapid-Fix, stained with Gill 3 hematoxylin and Eosin-Y, dehydrated with 95% and 100% ethanol, and cleared in xylene.  Cover slips were mounted with Shandon Mounting Media (Thermofisher Scientific, Kalamazoo, MI).  10X bright field images of each biopsy were captured with an Olympus BX61 microscope utilizing Cellsens Dimension^TM^ software. Image Pro Premier software (Mediacybernetics, Rockville, MD) was used to measure epidermal thickness, rete ridge path length and stratum corneum thickness in each biopsy.  The thickness of the viable epidermal was measured by tracing two separate lines along the dermal epidermal junction (DEJ) and the epidermal granular layer, then calculating the average distance between these two lines across the entire epidermal length.  Rete ridge path length was measured by taking the ratio of the DEJ length to that of the granular layer.  Stratum corneum thickness was measured on 20X bright field images by tracing two separate lines along both the epidermal granular layer and the top of the stratum corneum, then calculating the average distance between these two lines across the entire epidermal length.

**S.3 Proteomics**

For proteomics analysis (n=5 each of 20’s and 60’s photo-exposed arm samples), epidermal LCM sections were solubilized in 25 μl 0.5M TEAB (triethylammonium bicarbonate) containing 0.1% Rapigest. A 2 μl aliquot is taken for AAA (Amino Acid Analysis), and to the remaining amount 2.5 μl of 50 μM TCEP is added to reduce the cysteine residues. 1.3 μl of 200 μM MMTS was added to block cysteines. The proteins mixture was digested with LysC (1:10 wt/wt ratio) for 4 hours at 37°C, followed by trypsin addition to the mixture for digestion overnight at 37°C. Digest were then acidified and desalted using a micro-spin C18 RP column. Eluted peptide mixture was dried and reconstituted in UPLC loading Buffer A (0.1% formic acid) and Labelfree quantitative LC-MS/MS was performed on an LTQ Orbitrap Elite (ThermoFisher Scientific) equipped with a Waters Symmetry® C18 (180 µm x 20 mm) trap column and a 1.7 µm, 75 µm x 250 mm nanoAcquity™ UPLC™ column (35ºC). Trapping was done using 99% Buffer A (100% water, 0.1% formic acid) and peptide separation was undertaken using a linear gradient of solvents A (0.1% formic acid in water) and B (0.075% formic acid in acetonitrile) over 210 minutes, at a flow rate of 300 nL/min. MS spectra was acquired in the Orbitrap using 1 microscan and a maximum injection time of 900 ms followed by three data dependant MS/MS acquisitions in the ion trap (with precursor ions threshold of >3000). The total cycle time for both MS and MS/MS acquisition was 2.4 seconds. Peaks targeted for MS/MS fragmentation by collision induced dissociation (CID) were first isolated with a 2 Da window followed by normalized collision energy of 35%. Dynamic exclusion was activated where former target ions were excluded for 30 seconds.

Feature extraction, chromatographic/spectral alignment, data filtering, and statistical analysis were performed using Non-linear Dynamics Progenesis LCMS software (Non-linear Dynamics, LLC). First, the raw data files were imported into the program. A sample run was chosen as a reference (usually at or near the middle of all runs in a set), and all other runs were automatically aligned to that run in order to minimize retention time (RT) variability between runs. No adjustments are necessary in the m/z dimension due to the high mass accuracy of the mass spectrometer (typically <3 ppm). All runs were selected for detection with an automatic detection limit. Features within RT ranges of 0–16 min and 102–120 min was filtered out, as were features with charge ≥ +8. A normalization factor was then calculated for each run to account for differences in sample loading between injections. The experimental design was set up to group multiple injections from each run. The algorithm then calculates and tabulates raw and normalized abundances, max fold change, and ANOVA p-values for each feature in the data set. The MS/MS collected for the experiment were filtered to exclude spectra with rank > 10 or isotope > 3 to ensure that the highest quality MS/MS spectral data are utilized for peptide assignments and subsequent protein ID. The remaining MS/MS were exported to an .mgf (Mascot generic file) for database searching. After the Mascot search, an .xml file of the results was created, and then imported into the Progenesis LCMS software, where search hits were assigned to corresponding features.

**S.4 Skin surface biomarker analysis**

Stratum corneum material was collected from each subject’s dorsal arm, cheek, and buttock sites. Two sets of three sequential D-Squame tape strip samples (standard sampling discs, 22-mm diameter; CuDerm Corp., Dallas, TX, USA) were collected from each site. Tapes 2 and 3 from an individual site were designated for IL-8 and IL-1RA/IL-1α ratio analysis and tapes 2 and 3 from a neighboring site were used for biochemical metabolite analysis. The tape samples were collected, stored at -80°C, and total protein and metabolites were extracted as previously described.^Kerr^ IL-1RA and IL-1α levels were measured by ELISA analysis as per manufacturer’s instructions (Bio-Plex Pro Human Cytokine IL-1RA and IL-1α kits, Bio-Rad, Hercules, CA, USA). IL-8 was quantified using Meso Scale Discovery (MSD) electrochemiluminescence V-Plex kit as per manufacturer’s instructions and normalized to the amount of soluble total protein as determined by BCA protein assay (BCA™ Protein Assay Kit, Pierce Biotechnology/Thermo Scientific, Rockford, IL, USA). For *cis*- and *trans*-urocanic acid metabolite profiling, individual tubes containing D-Squame strips were pooled to generate 120 samples with 5 replicates per each collection site from 20’s, 40’s, and 60’s age groups. Metabolomic profiling analysis was performed by Metabolon Inc. (Durham, NC, USA) as previously described.^1^ Welch’s two-sample *t*-test, matched pair *t*-test, and Principal Component Analysis (PCA) were used to analyze the data.

^1^Evans AM, DeHaven CD, Barrett T, Mitchell M, Milgram E. Integrated, nontargeted ultrahigh performance liquid chromatography/electrospray ionization tandem mass spectrometry platform for the identification and relative quantification of the small-molecule complement of biological systems. *Analytical chemistry* 2009: **81**: 6656–6667.

**S.5 Epigenetic Aging clock (DNAge^®^)**

Skin DNA was bisulfite converted using the EZ DNA Methylation-Lightning™ Kit (Zymo Research, Irvine, CA, USA) according to the manufacturer’s instructions. Bisulfite-converted DNA libraries contains >2,000 age-associated CpG loci were prepared for Simplified Whole-panel Amplification Reaction Method (SWARM**^®^**) platform, which is a targeted bisulfite-based approach where specific CpG loci were sequenced at >1,000x coverage on a HiSeq sequencer. Sequence reads were identified by base calling software then aligned to the hg19 genome using Bismark, an aligner optimized for bisulfited converted sequences. Methylation levels for each cytosine were calculated by dividing the number of reads reporting a “C” by the number of reads reporting a “C” or “T.” The methylation level of >2,000 age-associated CpG loci were used for age prediction using Zymo Research’s proprietary DNAge**^®^** predictor.

**S.6 Immunofluorescence and immunohistology**

**CDKN2A/p16^INK4a^**

7 μm fresh frozen cryosections were fixed in ice cold acetone for 10 minutes at -20°C, washed in phosphate-buffered saline (PBS), and incubated for 1 hour at room temperature (RT) in 10% normal goat serum in PBS (Cell Signaling Tech, Danvers, MA 5425S).  Sections were incubated 1 hour at RT with an anti-CDKN2A/p16^INK4a^ (Abcam, Waltham, MA, ab108349 1:500) antibody, washed in PBS, incubated with an Alexa Fluor 555-conjugated goat anti-rabbit antibody (Abcam, Waltham, MA, ab150086 1:1000) for 1 hour at RT, washed in PBS and counterstained with DAPI using NucBlue fixed cell stain Ready Probes reagent (Invitrogen, Carlsbad, CA).  Staining of sections minus the primary antibody served as a negative control and displayed no non-specific staining (data not shown). For comparison fluorescent images of young and old biopsies were captured with a Zeiss Observer.Z1 microscope (Carl Zeiss Microimaging, Germany) at equal gamma values, pixel range and exposures.

**HIF-1α**

7 μm fresh frozen cryosections were fixed in ice cold acetone for 10 minutes at -20°C, washed in PBS, and incubated for 1 hour at RT in 10% normal goat serum in PBS (Cell Signaling Tech, Danvers, MA 5425S).  Sections were incubated overnight at 4°C with an anti-HIF-1α antibody (Sigma, ST. Louis, MO, HPA001275 1:100), washed in PBS, incubated with an Alexa Fluor 555-conjugated goat anti-mouse antibody (Abcam, Waltham, MA, ab150118 1:1000) for 1 hour at RT, washed in PBS, and counterstained with DAPI using NucBlue fixed cell stain Ready Probes reagent (Invitrogen Carlsbad, CA).  Staining of sections minus the primary antibody severed as a negative control and displayed no non-specific staining (data not shown). For comparison fluorescent images of young and old biopsies were captured with a Zeiss Observer.Z1 microscope (Carl Zeiss Microimaging, Germany) at equal gamma values, pixel range and exposures.

**Hemoglobin-α**

7 μm fresh frozen cryosections were fixed in ice cold Acetone for 10 minutes at -20°C, washed in (PBS), and incubated for 30 minutes at RT in 5% normal mouse serum in PBS (Invitrogen Carlsbad, CA, 31881).  Sections were incubated 90 minutes at RT with an anti-hemoglobin-α antibody (Santa Cruz Biotech, Dallas, TX, sc-514378 AF488 1:100), washed in PBS, counterstained using NucBlue fixed cell stain Ready Probes reagent (Invitrogen CA, U.S.A.).  Mouse IgG1 isotype negative controls (Invitrogen Carlsbad, CA MA5-18167 1:100) displayed no non-specific staining. For comparison fluorescent images of young and old biopsies were captured with a Zeiss Observer.Z1 microscope (Carl Zeiss Microimaging, Germany) at equal gamma values, pixel range and exposures.

**Blood vessel staining utilizing *Ulex Europaeus*-I Lectin (UEA-1)**

7 μm fresh frozen cryosections were fixed in 95% ethanol at RT for 2 minutes, washed in PBS, and incubated with FITC conjugated UEA-1 (Sigma, St. Louis, MO, L9006 1:50 in water) for 2 minutes at RT, washed in PBS, and coverslipped using Fluorshield with DAPI (Sigma, MO, U.S.A., F6057). For comparison fluorescent images of young and old biopsies were captured with a Zeiss Observer.Z1 microscope (Carl Zeiss Microimaging, Germany) at equal gamma values, pixel range and exposures.

**53BP1**

Frozen skin sections were briefly thawed and circled with a hydrophobic barrier pen. Sections were rehydrated with PBS and were fixed with 3% paraformaldehyde for 15 minutes at RT. After 2 washes in PBS, sections were subsequently permeabilized with 0.5% Triton-X for 10 minutes at RT. After 3 washes in PBS, sections were blocked with 5% normal donkey serum for 30 minutes at RT. Subsequently, they were incubated overnight in primary antibodies (1:1000 anti-53BP1 Novus Biologicals and 1:250 anti-K10 Dako in 5% normal donkey serum) at 4°C in a humidified chamber. After 3 washes in PBS, appropriate fluorophore-conjugated secondary antibodies (1:800 donkey anti-rabbit Alexa Fluor 564, 1:800 donkey anti-mouse Alexa Fluor 488 in 5% normal donkey serum) were added for 1 hour at RT. Hoechst dye was used as a nuclear counter-stain. After 3 washes in PBS, sections were mounted in Prolong-Diamond Anti-Fade reagent. Imaging was done at 60X magnification on an Olympus IX-83 inverted fluorescence microscope. Z-stacks were acquired for each position along the length of the skin sample. Projection images were made for each position using Image J and the number of cells showing visible 53BP1 foci were quantified in the K10-positive suprabasal layer and K10-negative basal layer.

**Filaggrin and Involucrin**

10 μm fresh frozen sections were fixed in ice cold acetone and methanol (1:1) for 10 minutes at -20°C, washed in PBS and incubated for 1 hour at RT in 10% normal goat serum in PBS (Cell Signaling Tech, Danvers, MA 5425S).  Sections were incubated overnight at 4°C with an anti-filaggrin (Abcam, Waltham, MA, ab3137 1:100) or an anti-involucrin (Sigma, St. Louis, MO, I9018 1:100) antibody, washed in PBS, incubated with Alexa Fluor 488 conjugated goat anti-mouse antibodies (Abcam, Waltham, MA, ab1500113 1:500) for 1 hour at RT, washed in PBS, and mounted with fluoroshield containing DAPI (Sigma, St. Louis, MO, F6057).  Mouse IgG1 isotype negative controls (Invitrogen Carlsbad, CA MA5-18167 1:100) displayed no non-specific staining (data not shown).  For comparison fluorescent images of young and old biopsies were captured with a Zeiss Observer.Z1 microscope (Carl Zeiss Microimaging, Germany) at equal gamma values, pixel range and exposures.

**Loricrin**

7 µm fresh frozen sections were fixed in ice-cold 50% acetone/50% methanol at RT for 5 minutes. Sections were air dried followed by 3 washes in PBS/0.05% Tween 20. They were then blocked with 10% goat serum in PBS for 30 minutes. Subsequently, sections were exposed to anti-loricrin antibody (1:500; Abcam, Singapore, Singapore, ab176322) for 1 hour followed by a 30-minute incubation with an anti-rabbit Alexa Fluor 568 (1:1000; Thermo Fisher, Singapore, Singapore) and a 10-minute counterstain with DAPI (Sigma–Aldrich, Singapore, Singapore) before mounting with Hydromount™ (Electron Microscopy Sciences).

**Keratin 14 and Keratin 10**

Skin biopsies were fixed in 4% paraformaldehyde (Sigma-Aldrich, Missouri, United States), serially dehydrated in ethanol, then incubated in Histo-Clear (Scientific Laboratory Supplies, Nottingham, United Kingdom) for 30 minutes, and a 1:1 ratio of Histo-Clear and paraffin wax (Thermo Fisher Scientific, Massachusetts, United States) for 60 minutes. Models were incubated in paraffin wax for 1 hour at 65^o^C prior to embedding (Solmedia Ltd, Shrewsbury, United Kingdom). 5 μm sections were generated using a microtome (Leica, Wetzlar, Germany) and transferred onto charged microscope slides (Thermo Fisher Scientific). Skin sections were deparaffinised in Histo-Clear and rehydrated from 100% ethanol to PBS. Antigen retrieval was performed using pH 6 citrate buffer at 95^o^C for 20 minutes. Samples were blocked and permeabilised for 1 hour in a blocking buffer of 20% neonatal calf serum (Thermo Fisher Scientific, Massachusetts, United States) in 0.4% Triton X-100 in PBS. Sections were incubated overnight in primary antibodies (1:100 cytokeratin 14 ab7800; 1:100 cytokeratin 10 ab76318; Abcam, Cambridge, UK) at 4^o^C in a humidified chamber. After 3 washes in PBS, appropriate fluorophore-conjugated secondary antibodies (1:1000 donkey anti-mouse Alexa Fluor^®^ 488, 1:1000 donkey anti-rabbit Alexa Fluor^®^ 594) were added for 1 hour at RT. After 3 washes in PBS, sections were mounted in Vectashield Hardset with DAPI mounting medium (Vector Laboratories, Peterborough, United Kingdom). 40X images were captured at equal gamma values, pixel range and exposures using a Zeiss 880 confocal microscope (Zeiss, Oberkochen, Germany) with Zen software.
