## Supplementary Table 1 for "Inflammaging in human photoexposed skin: Early onset of senescence and imbalanced epidermal homeostasis across the decades"

Table 1. Quantification of histological measurements of epidermal morphology across age groups and face, arm, and buttock sites (mean ± SEM).

| Body site | Age group | Replicates | Stratum corneum thickness (μm) ± SEM | Epidermal thickness (μm) ± SEM | Rete ridge length ratio  ± SEM |
| --- | --- | --- | --- | --- | --- |
| *Face* | 20’s | 30 | 10.69 ± 0.52 | 66.67 ± 1.80 | 1.220 ± 0.025 |
|  | 30’s | 24 | 10.82 ± 0.47 | 63.25 ± 2.12 | 1.210 ± 0.032 |
|  | 40’s | 24 | 13.35 ± 0.84* | 64.95 ± 2.25 | 1.242 ± 0.047 |
|  | 50’s | 26 | 13.31 ± 0.63** | 60.18 ± 1.88* | 1.205 ± 0.030 |
|  | 60’s | 22 | 13.45 ± 1.04* | 56.95 ± 2.71** | 1.172 ± 0.043 |
|  | 70’s | 26 | 14.44 ± 0.71*** | 51.92 ± 1.83*** | 1.086 ± 0.025** |
| *Arm* | 20’s | 29 | 21.98 ± 0.90 | 57.45 ± 1.76 | 1.182 ± 0.029 |
|  | 30’s | 25 | 24.30 ± 1.52 | 56.80 ± 1.70 | 1.125 ± 0.027 |
|  | 40’s | 24 | 22.94 ± 1.14 | 52.84 ± 1.48 | 1.119 ± 0.027 |
|  | 50’s | 26 | 25.45 ± 1.50 | 55.41 ± 2.86 | 1.091 ± 0.027* |
|  | 60’s | 22 | 24.17 ± 1.20 | 46.25 ± 2.50* | 1.098 ± 0.018 |
|  | 70’s | 23 | 25.74 ± 1.94 | 50.87 ± 2.31* | 1.057 ± 0.019* |
| *Buttock* | 20’s | 27 | 17.83 ± 0.69 | 74.13 ± 2.99 | 1.449 ± 0.042 |
|  | 30’s | 25 | 19.97 ± 0.97 | 67.86 ± 1.76 | 1.464 ± 0.057 |
|  | 40’s | 24 | 20.46 ± 0.78* | 72.70 ± 3.33 | 1.559 ± 0.049 |
|  | 50’s | 26 | 19.10 ± 0.87 | 61.01 ± 2.04*** | 1.446 ± 0.049 |
|  | 60’s | 22 | 21.32 ± 1.31* | 59.91 ± 2.35*** | 1.398 ± 0.049 |
|  | 70’s | 25 | 21.23 ± 0.94** | 58.91 ± 2.07*** | 1.300 ± 0.036* |

T-test (two tailed, type III) comparing each decade to the 20’s age group. * *p*<0.05; ** *p*<0.01; *** *p*<0.001
